# Tethered homing gene drives: a new design for spatially restricted population replacement and suppression

**DOI:** 10.1101/457564

**Authors:** Sumit Dhole, Alun L. Lloyd, Fred Gould

## Abstract

Optimism regarding potential epidemiological and conservation applications of modern gene drives is tempered by concern about the potential unintended spread of engineered organisms beyond the target population. In response, several novel gene drive approaches have been proposed that can, under certain conditions, locally alter characteristics of a population. One challenge for these gene drives is the difficulty of achieving high levels of localized population suppression without very large releases in face of gene flow. We present a new gene drive system, Tethered Homing (TH), with improved capacity for localized population alteration, especially for population suppression. The TH drive is based on driving a payload gene using a homing construct that is anchored to a spatially restricted gene drive. We use a proof of principle mathematical model to show the dynamics of a TH drive that uses engineered underdominance as an anchor. This system is composed of a split homing drive and a two-locus engineered underdominance drive linked to one part of the split drive (the Cas endonuclease). In addition to improved localization, the TH system offers the ability to gradually adjust the genetic load in a population after the initial alteration, with minimal additional release effort.

## INTRODUCTION

The development of population genetic theory related to use of translocations and other underdominance mechanisms to suppress pest populations or change their characteristics started more than five decades ago (Serebrovskii 1940, Vanderplank 1947, Curtis 1968). Under the best of circumstances these approaches were expected to require release of large numbers of genetically manipulated individuals, and would have only localized impacts. Despite major efforts, early empirical work using these approaches were unsuccessful (Gould and Schliekelman 2004). Advances in transgenic techniques for engineering insects spurred the hope that natural populations of insect pests could be suppressed or manipulated using transposable elements (Ribeiro and Kidwell 1994, O’Brochta et al. 2003) and other types of selfish genetic elements (Sinkins and Gould 2006). The expectation was that fewer individuals would need to be released and the spread would not be localized (Burt 2003). In spite of the slow initial progress on such approaches (Carareto et al. 1997, Chen et al. 2007, Windbichler et al. 2008), researchers and popular media commentators raised concerns that unrestricted spread could be problematic and that safeguards were needed (Burt 2003, Gould 2008). With the recent development of CRISPR/Cas-based gene drives (Esvelt et al. 2014, Gantz et al. 2015), such concerns have intensified, and refocused attention on spatially and temporally restricted gene drives as safer alternatives for many applications (e.g. Akbari et al. 2015, NASEM 2016, Min et al. 2017, Marshall and Akbari 2018).

A number of strategies have been put forth for engineered gene drives that are expected to be relatively restricted either spatially (Davis et al. 2001, Marshall and Hay 2012a, Akbari et al. 2014, Buchman et al. 2018), temporally (Gould et al. 2008) or both temporally and spatially (Rasgon 2009, Noble et al. 2016, Burt and Deredec 2018). Marshall and Hay (2012a) and Dhole et al. (2018) have compared some of the properties of these gene drive systems using simple population genetics models. The release size required for different gene drives to successfully invade a population varies widely even when the drive constructs do not impose high fitness costs. The spatially restricted drives are generally not expected to establish themselves at high frequency in neighboring populations when migration rates are low (< 1%; but see some results in Dhole et al. 2018, Champer et al. 2018a). As migration rates to neighboring populations increase, spatial restriction to the targeted population is not assured (Marshall and Hay 2012a, Dhole et al. 2018, Champer et al. 2018a).

If instead of population replacement, the goal of a release is population suppression, the drive constructs must impose high fitness costs, and because of that, the spatially confined drives require much larger releases to spread into a local population, if they can spread at all (Magori and Gould 2006, Ward et al. 2011, Marshall and Hay 2012a, Dhole et al. 2018, Edgington and Alphey 2018, Khamis et al. 2018). In general, gene drives that require larger releases when the constructs have low fitness costs tend to remain more localized to the target population. However, these gene drives are also the least likely to be able to drive constructs with high fitness costs into the target population (Marshall and Hay 2012a, Dhole et al. 2018).

There is a need for a gene drive that is reasonably confined and can spread constructs with high fitness costs. We recently described a spatially restricted gene drive for population suppression that relies on a CRISPR/Cas endonuclease that disrupts an allele that is fixed in the target population, but cannot disrupt other alleles at the same locus that are found at least at low frequencies in neighboring populations (Sudweeks et al. in review). That approach would be specifically appropriate for small populations on oceanic islands where genetic drift is expected to be strong. Min et al. (2017) have also outlined a verbal model of a gene drive construct, the daisy quorum drive, that may potentially allow localized spread of high cost payloads with relatively small release size. A mathematical exploration of the dynamics of the daisy quorum drive is not yet available.

Here we propose a new concept of Tethered Homing (TH) gene drives. These drives include a homing component that does not drive on its own, but is “tethered” by engineering it to be reliant on a spatially restricted gene drive. We present the dynamics of a specific TH gene drive design that uses a two-locus engineered underdominance component to tether a CRISPR-Cas-based homing component (Underdominance Tethered Homing, UTH). Conceptually, the homing component can instead be tethered to a different localized gene drive, such as one-locus engineered underdominance, chromosomal translocations, or one of the poison-antidote systems (Marshall and Hay 2012b, Akbari et al. 2013).

As with other analyses aimed at initial description of novel gene drive systems (Davis et al. 2001, Gould et al. 2008, Marshall and Hay 2012b, Burt and Deredec 2018), we explore the properties of the UTH strategy using a very general, proof-of-principle mathematical model (Servedio et al. 2014). In addition to describing the general dynamics of the UTH drive, our analyses are specifically intended to facilitate direct comparison of this drive with three previous gene drives designed for localized population alteration – the daisy chain drive (Noble et al. 2016), and two engineered underdominance drives (Davis et al. 2001). We demonstrate that a UTH gene drive can offer an improvement in localization level. A UTH drive can especially be useful to locally spread high-cost payload genes while having relatively small effects on neighboring populations.

### Drive design

The underdominance tethered homing (UTH) drive is a two-component drive. The first component is a two-locus engineered underdominance drive linked to genes for producing a Cas endonuclease. The second component is a construct containing sequences coding for multiple guide RNAs that, in the presence of the Cas endonuclease, target the wild-type gene on the homologous chromosome, in turn triggering homing through the cell’s homology directed repair (HDR) pathway (see details below).

The two-locus underdominance component is structured following the design proposed by Davis et al. (2001), with the addition of sequence for germline-specific production of a Cas endonuclease. It is composed of two constructs (Figure 1A), located at separate loci. In our model, the two loci are termed A and B, with the transgenic alleles labeled A_*t*_ and B_*t*_. The corresponding wild-type alleles at the loci are referred to as A_*w*_ and B_*w*_. Each underdominance construct is engineered to produce a lethal toxin during early life stages, unless the individual also possesses a copy of the other underdominance construct, which harbors a suppressor for transcription of the toxin gene on the first construct (Figure 1A). Thus, only wildtype individuals and those that carry at least one copy of each transgenic allele (A_*t*_ and B_*t*_) are viable.

**Figure 1:**
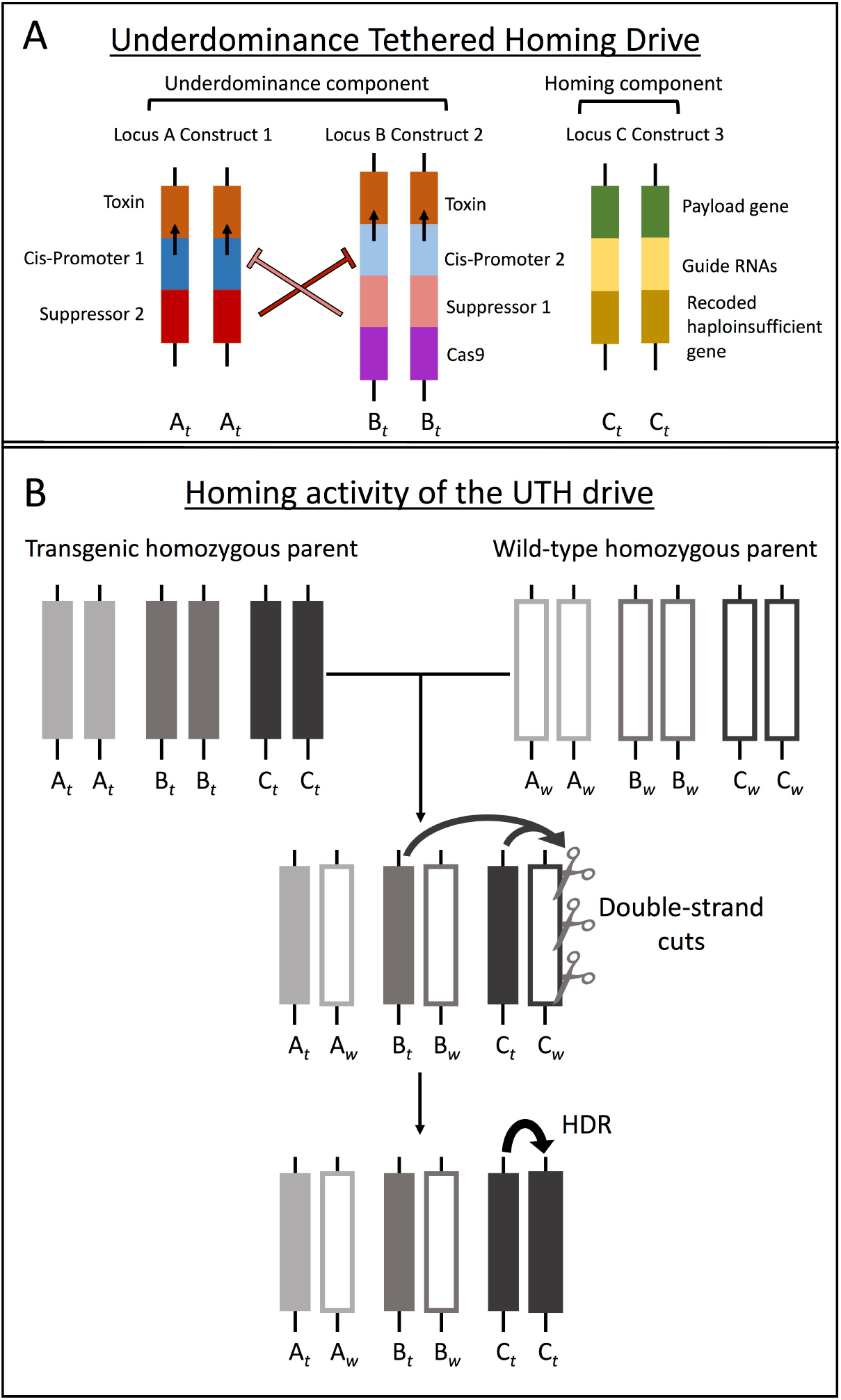
The general design and mechanism of the underdominance tethered homing (UTH) drive are shown. Different genetic elements that form a construct are shown as colored segments. A) The three constructs of the UTH drive occupy three unlinked loci. The two underdominance constructs form a toxin-suppressor engineered underdominance system. The third construct forms a homing component that is driven in presence of the other two constructs. B) In the germline of heterozygous individuals, the Cas endonuclease from the underdominance component along with the guide RNAs target the wild-type counterpart of the homing construct for multiple cuts. Repair through the cell’s homology directed repair (HDR) pathway leads to insertion of the homing construct into the homologous chromosome, rendering the cell homozygous for the homing construct.

Only engineered individuals that carry both underdominance constructs are viable, therefore the sequence for the Cas endonuclease does not need to be included in both underdominance constructs. In the model shown here it is included only in allele B_t_. Accidental nuclease activity of Cas in absence of any guide RNAs or even the resource cost of producing the Cas protein may impart some fitness cost to the underdominance component. We therefore assume that the allele *B_t_* incurs a multiplicative fitness cost. The parameter s_*c*_ gives the cost paid by individuals homozygous for the B_*t*_ allele. Results with equal cost for both underdominance constructs are included in the supplementary material, and are qualitatively similar.

The homing component of the UTH drive is located at a third, unlinked locus ‘C’. This component is specifically designed to target and be inserted into a haploinsufficient gene, i.e. two copies of a functional gene are required at this locus for embryonic development or for gametogenesis. Engineered and wild-type alleles at this locus are denoted C_*t*_ and C_*w*_, respectively The homing component is composed of three tightly linked segments: 1) sequences for multiple guide-RNAs with promoters that target the wild type haploinsufficient gene (C_*w*_), 2) a modified copy of the wild-type gene that cannot be targeted by the guide RNAs, and 3) a payload gene with a promoter suited for the payload’s function (Figure 1A). The homing construct is assumed to incur a multiplicative fitness cost (due to the payload and guide RNA production), where the cost paid by C_*t*_ homozygotes is given by parameter *s_p_*.

In heterozygous germline cells, the guide RNAs and the Cas endonuclease together target the wild-type (C_*w)*_ alleles for multiple double-stranded breaks (Figure 1B). If the damage is repaired through fully successful HDR, this results in germline cells that are homozygous for the homing construct. Repair through nonhomologous end-joining (NHEJ) would be expected to result in a deletion at this locus due to the multiple breaks. Nonhomologous end-joining or HDR that did not produce a functional copy of the haploinsufficient gene would be expected to result in individuals that are incapable of producing viable offspring (due to the haploinsufficiency at the locus). These two design elements, multiplexed guide RNAs and a haploinsufficient target gene, are expected to prevent (or drastically reduce) the emergence of alleles resistant to the drive (Esvelt et al. 2014; Noble et al. 2016; Noble et al. 2017). For this reason we assume no resistance alleles in our model. The homing efficiency (*H*) of the drive describes the likelihood of successful HDR at all sites identified for cutting by the guide RNAs. Thus, after the action of Cas endonuclease and the guide RNAs an individual’s fitness is reduced, on average, by a factor of (1-*H*) due to inefficient homing, in addition to fitness reduction due to the costs of the two drive components (*s_c_* and *s_p_*). The relative fitness of all genotypes is given in Table S1 in the supplementary material.

### Two-genotype release

When the UTH drive is introduced into a population, the release group is composed of individuals of two genotypes. The majority of the individuals released carry only the underdominance component (genotype A_*t*_A_*t*_B_*t*_B_*t*_C_*w*_C_*w*_), and only a small fraction of the individuals released also include the homing component that includes the payload gene (genotype A_*t*_A_*t*_B_*t*_B_*t*_C_*t*_C_*t*_). The underdominance constructs are expected to increase rapidly in frequency in a population if introduced above a threshold frequency (Davis et al. 2001; Edgington and Alphey 2017; Dhole et al. 2018). The lower release frequency of the homing component is intended to prevent it from imposing high indirect selection (i.e. selection due to linkage disequilibrium) against the underdominance component before underdominance reaches a high frequency (see Results).

### Alternative two-stage delayed release

Instead of a single two-genotype release, it is also possible to introduce the UTH drive with a two-stage delayed release, where the releases of individuals of different genotypes are separated temporally. For such an introduction, only individuals carrying the underdominance component are released initially. Individuals homozygous for the whole UTH drive are then release after a delay of 10 generations. This release scheme also allows underdominance to establish in the population by delaying the burden imposed by the cost of the payload gene.

### Simulations

We use population genetic simulations to describe the dynamics of the UTH gene drive in large, well-mixed populations. We assume a 1:1 sex ratio at birth and random mating based on adult genotypic frequencies. We also assume that all drive-based natural selection (fitness impacts of constructs, segregational cost of underdominance and inefficient homing) occurs in pre-adult stages, which may reduce the size of the adult population within a generation, but we assume that the same number of offspring are born every generation. This assumes perfect density compensation. We first address drive dynamics in a single population, and then use a two-population setting to study the level of localization for the UTH gene drive.

Two factors that strongly influence the performance of a homing-based gene drive are the fitness costs of the drive components (here given by parameters *s_c_* and *s_p_*), and the homing efficiency (given by parameter *H*). The underdominance component of the UTH drive only serves the purpose of providing the Cas endonuclease for the homing construct. Therefore, the same underdominance constructs can be used for substantially different applications of the gene drive. The costs of the payload gene, on the other hand, will vary depending upon the purpose of the gene drive release. Homing efficiencies for CRISPR/Cas-based homing drives vary across species and across genetic constructs (Champer et al. 2018b). We analyze the release effort required to alter an isolated population with a UTH drive under a range of possible values for the cost of the underdominance component, for payload costs, and for the homing efficiency.

The results shown below are from simulations of a single two-genotype release, with the exception of two cases that are highlighted. We simulate a release where engineered individuals of both sexes are introduced. The starting frequency, after the single initial release, for the homing construct is 5%, while the frequency of the underdominance component varies from 5% to 90%. As a measure of population alteration, we use the mean frequency of the gene drive constructs over a 100 generation period after the release. The mean frequency gives a more accurate representation of population alteration than the final allelic frequencies.

### Capacity for population suppression

It is clear that a gene drive intended for population suppression must impose a high genetic load on the target population (i.e. reduction in the mean fitness or reproductive capacity of individuals in the altered population compared to a population composed entirely of wildtype individuals) (Burt 2003). Gene drives that can impose high genetic loads, for sustained periods, are likely to achieve substantial population suppression (Burt 2003, Sinkins and Gould 2006). A rigorous analysis of population suppression would require a model that can incorporate biological details much beyond the scope of our proof-of-principle model. Details of a population’s density-dependent dynamics, mating behavior as well as stochasticity are expected to play major roles in determining how a population responds to genetic load imposed by any gene drive (e.g. Edgington and Alphey 2018, Khamis et al. 2018). For many species, spatial dynamics within a population will also likely influence gene drive dynamics and the level of population suppression. Migration between multiple populations can further complicate suppression analysis. These population-specific details will need to be incorporated in future models.

To gain a comparative measure of the capacity of the UTH drive for population suppression, we use the level and duration of genetic load it can impose, and compare this with similar measures for previously proposed gene drives. We show the mean genetic load imposed by the UTH drive on an isolated population after single two-genotype releases of varying sizes. Reducing female viability (or fecundity) has a stronger influence on population genetic load than reducing male viability (or fertility). Suppression drives that only reduce female fitness are also likely to be more efficient, because natural selection acting against the drive would be weak in males. Also, a payload with recessive fitness costs is expected to face weaker natural selection when at low frequencies, facilitating easier initial spread (Magori and Gould 2006). For these reasons we model a UTH drive with a payload gene that only reduces female fitness by imposing a recessive fitness cost. Results with multiplicative fitness costs are provided in the supplementary material (Figure S2).

### Gradually increasing the genetic load

For certain applications, it may be desirable to be able to gradually increase or adjust the genetic load in a population. A useful feature of the UTH drive is that it can be used to successively drive multiple payload genes into a population with minimal release effort after the initial establishment of the drive. The additional payloads would be released as individuals with separate homing constructs (transgenic alleles at new loci D, E and so on) linked with new guide RNAs that would use the pre-established Cas gene. We simulate the spread of three payload genes designed for reducing female fitness, successively released at 20-generation intervals.

### Migration and localization analysis

To assess the level of localization of the UTH drive, we use a scenario with two populations, a Target and a Neighbor population, that exchange migrants every generation. We assume that the two populations are initially of equal size. However, the spread of the gene drive may alter population size within a generation if the gene drive constructs are costly. We assume that each adult individual has a fixed probability of migrating out of its native population before mating. Thus a constant fraction, *μ*, of individual from each population leave to join the other population. This means that if the number of adults in one of the population becomes smaller than the other, a smaller absolute number of individuals migrate out of it, relative to those migrating out of the larger population (e.g. Dhole et al. 2018). This is more realistic than fixed migration rates that imply the same number of individuals migrate irrespective of a population’s size. We term the effective immigration rate into the target population as *μ_T_*, and that into the neighbor population as *μ_N_*. We account for the potential differences in effective immigration rates (due to differences that may arise in adult population size) with the approximations

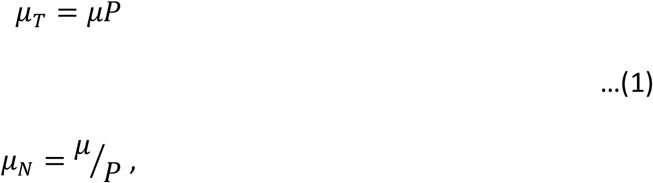

where *P* is a proxy for the ratio of the population sizes, given by

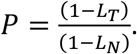

Here *L_T_* and *L_N_* are the genetic loads in a given generation in the target and the neighboring populations, respectively. Equations (1) allow for an asymmetry to arise in the effective immigration rates if the two populations begin to differ in size (when *P* ≠ 1). If both populations are of equal size (when *P* = 1), migration would be symmetrical. Note that the equations (1) do not incorporate changes in population size over multiple generations, but only within a generation. Explicitly modeling across-generation changes in population size would require a model that can incorporate population density and spatial dynamics, which are beyond the scope of this model. Analyses without any correction for differences in population size are included in the supplementary material.

We follow the frequencies of the three gene drive constructs (the three transgenic alleles at the three loci) in both populations though 100 generations after the initial release. Localized spread of the gene drive would result in successful alteration of the target population, while leaving the neighbor population largely unaltered.

## RESULTS

### Dynamics in an isolated population

The two-genotype release is a critical feature needed for efficient use of the UTH drive. Introducing the UTH drive into a population as a one-genotype release of individuals homozygous for the complete drive (genotype A_*t*_A_*t*_B_*t*_B_*t*_C_*t*_C_*t*_) results in drive failure, unless the release is extremely large or all drive constructs, including the payload, have extremely low fitness costs. For instance, a UTH drive carrying a payload with a 50% homozygous fitness cost fails after a single one-genotype introduction at 1:1 release ratio with wild-type (Figure 2A). However, a two-genotype release of the same size (1:1 release giving a starting frequency of 0.5 for alleles A_*t*_ and B_*t*_, and 0.05 for allele C_*t*_) results in successful drive (Figure 2B). The reason for the major difference between the success of these release approaches is that the cost of the homing construct (which carries the payload) results in strong selection against all components of the UTH drive, because strong linkage disequilibrium develops between the three loci upon release. The low initial frequency of the homing construct in a two-genotype release lets underdominance reach a high frequency, which then successfully drives the homing component to a high frequency. A two-stage delayed release of the homing component can similarly allow successful drive (Figure 2C).

**Figure 2:**
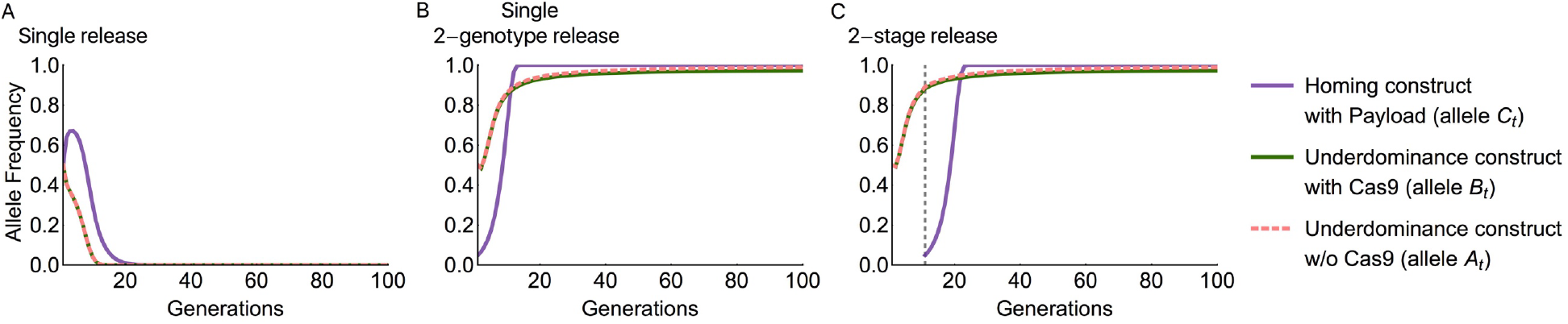
Time-series of different release methods for the UTH drive are shown, after a release of engineered individuals at a 1:1 ratio to wild-type individuals. Homing efficiency, *H*=0.95; underdominance construct cost for B_*t*_B_*t*_ homozygotes, *s_c_*=0.05; homozygous payload cost, *s_p_*=0.5.

The underdominance component causes the UTH drive to have a threshold release frequency that needs to be exceeded for successful population alteration. As expected for any case of underdominance (Sinkins and Gould 2006, Magori and Gould 2006, Altrock et al. 2010, Reeves et al. 2014, Edgington and Alphey 2017), higher costs of the underdominance components result in higher threshold frequencies (Figure 3 top row). Even when the release exceeds the threshold, higher costs of the underdominance component lower the equilibrium frequency of the underdominance constructs of the UTH drive in the population as indicated by the lighter red color representing the mean frequency over 100 generations. However, note that even these lower equilibrium frequencies of the underdominance constructs are sufficient to successfully drive the payload gene on the homing construct to high frequencies (Figure 3 bottom row). When the drive is released below the threshold, population alteration fails completely as none of the drive components spread (indicated by the dark blue color in the top and bottom rows of Figure 3). When the payload cost is too high, the drive fails as the homing construct is lost and only the underdominance component becomes established in the population (Figure 3; red area in top panel with corresponding blue area in bottom panel).

**Figure 3:**
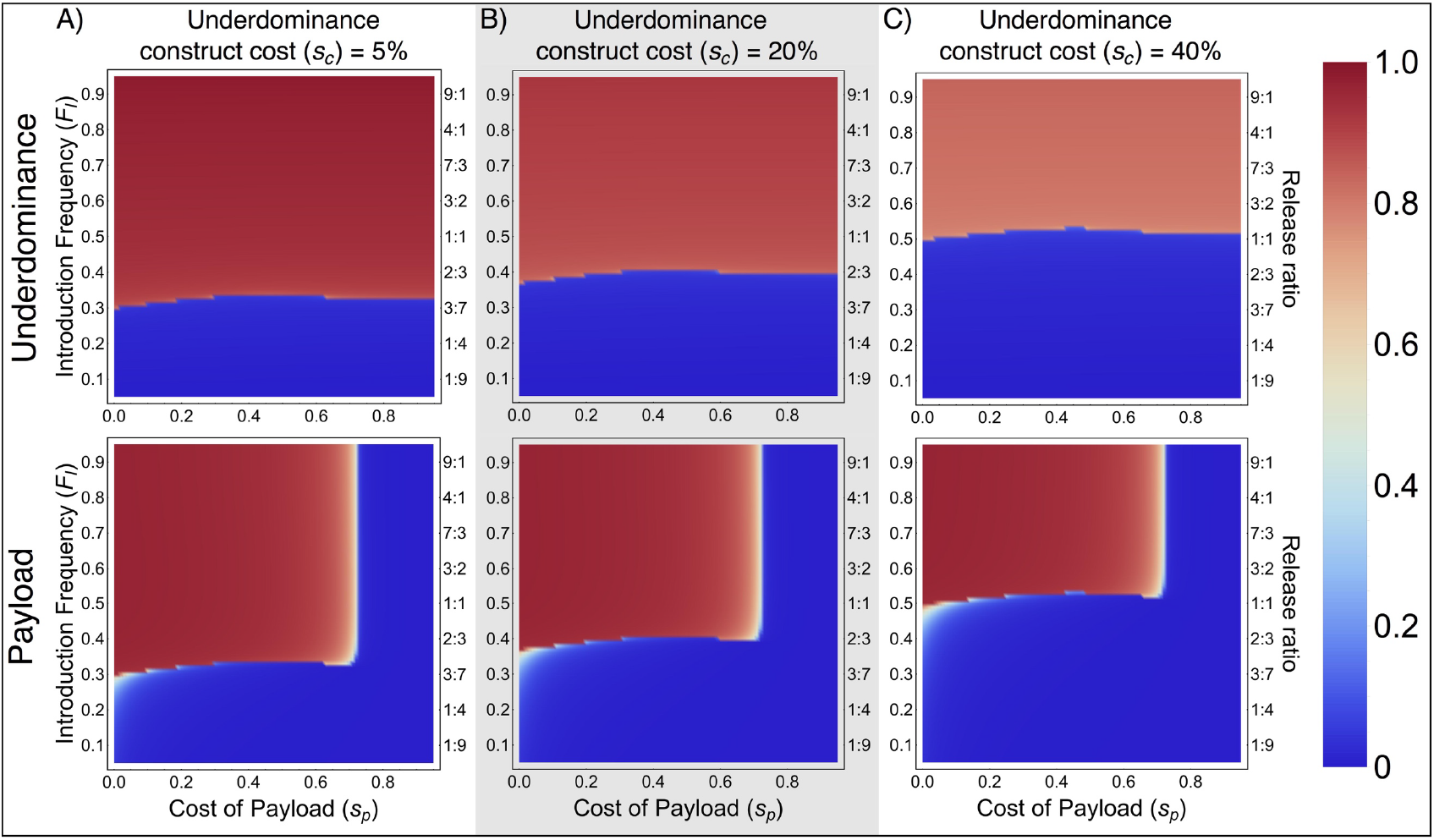
UTH drive in an isolated population - Colors show mean allelic frequencies for one of the underdominance (allele B_*t*_, top row) and the payload gene (bottom row) over a 100 generation time span following a single release. The three columns show results for UTH drives with different fitness costs of the underdominance component.

The relative ability of the UTH drive to push a payload gene into an isolated population can be compared with that of three other gene drives for which we have performed identical analyses – One- and two-locus engineered underdominance drives and the daisy chain drive (Dhole et al. 2018). Mean payload frequencies achieved with the UTH drive when assuming a 5% fitness cost of the underdominance (bottom panel of figure 3a) can be compared to figure 3 of Dhole et al. (2018). In the simple engineered underdominance drives analyzed in Dhole et al. (2018) the payload is directly linked to one of the underdominance constructs. This comparison shows that when the payload has high fitness costs, the UTH drive is dramatically more efficient than the simple engineered underdominance drives. The UTH drive can drive a payload gene with a given cost to a much higher frequency and with smaller release than can be done with a simple engineered underdominance drive. The daisy chain drive requires the lowest release, but has other drawbacks compared to the UTH drive (see results on localization in Dhole et al. 2018).

Results shown in Figure 3 are for simulations with 95% homing efficiency and equal payload fitness costs to both sexes. Lower homing efficiencies restrict the maximum payload gene cost that can still allow successful population alteration. Although, even with homing efficiencies as low as 70%, the UTH drive can successfully spread payloads with almost 50% homozygous fitness cost to both sexes, although the gene drive requires a large number of generations for complete population alteration (see supplementary material, Figure S3).

### Capacity for population suppression

A UTH drive designed for population suppression through high female-limited fitness costs can impose a considerable genetic load on a population (Figure 4). The underdominance component of the UTH drive imparts some genetic load due to the direct fitness cost of the constructs (*s_c_*) and the segregational cost of underdominance itself (lower fitness of heterozygotes). However, a large fraction of the genetic load is accrued through the spread of a costly payload located on the homing construct (*s_p_*). As the homing construct of the UTH is released at a low frequency, the genetic load builds up slowly. The mean genetic load in the first 20 generations after releasing the drive is therefore lower than the genetic load imposed in the 20^th^ generation – “final load” (Figure 4). For example, even with a very large release, the highest mean genetic load in the first twenty generations remains below 0.8, while the genetic load in the 20th generation can reach very close to unity for very costly payloads.

**Figure 4:**
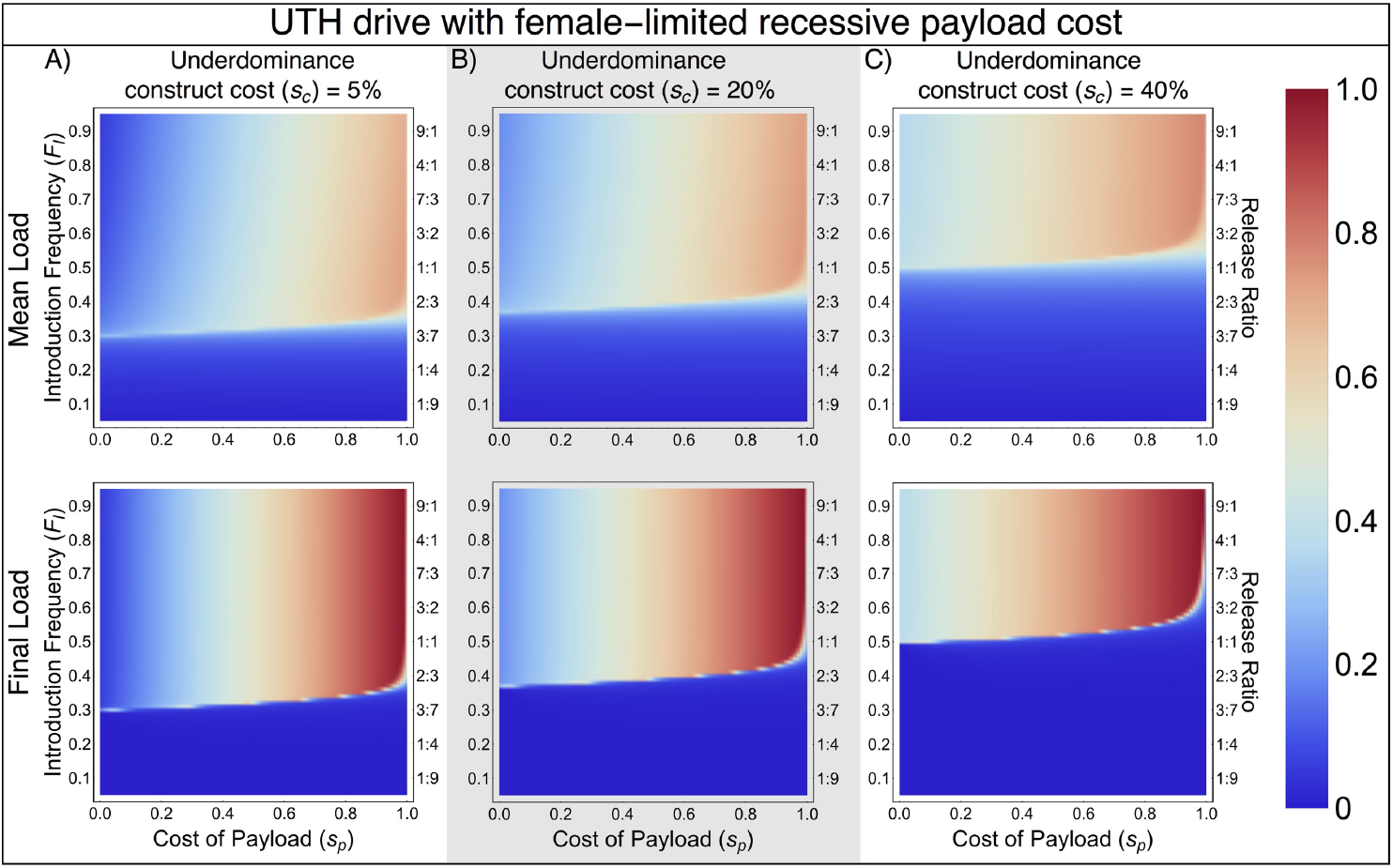
Colors show mean genetic load over twenty generations after drive release (top row) and final genetic load in the twentieth generation (bottom row).

With successive release of multiple payloads on different homing components, the UTH drive can be used to gradually adjust (increase or decrease) the genetic load imposed on a population. For example, a successive release of three payload genes with female-limited fitness reduction can be used to gradually increase the genetic load on a population to a high level (Figure 5). Conversely, a “rescue” payload gene can be driven to reduce the genetic load imposed by a loss of function payload gene. The total number of transgenic individuals released to spread the three payloads is only slightly higher than that for spreading a single payload, because after the initial establishment of the underdominance constructs, each additional payload only needs to be released at very low frequency (1% for the example shown in figure 5). The UTH drive can achieve high genetic loads using a broader set of parameters than with the other drives examined in Dhole et al. (2018—see figures 6 & S11 therein).

**Figure 5:**
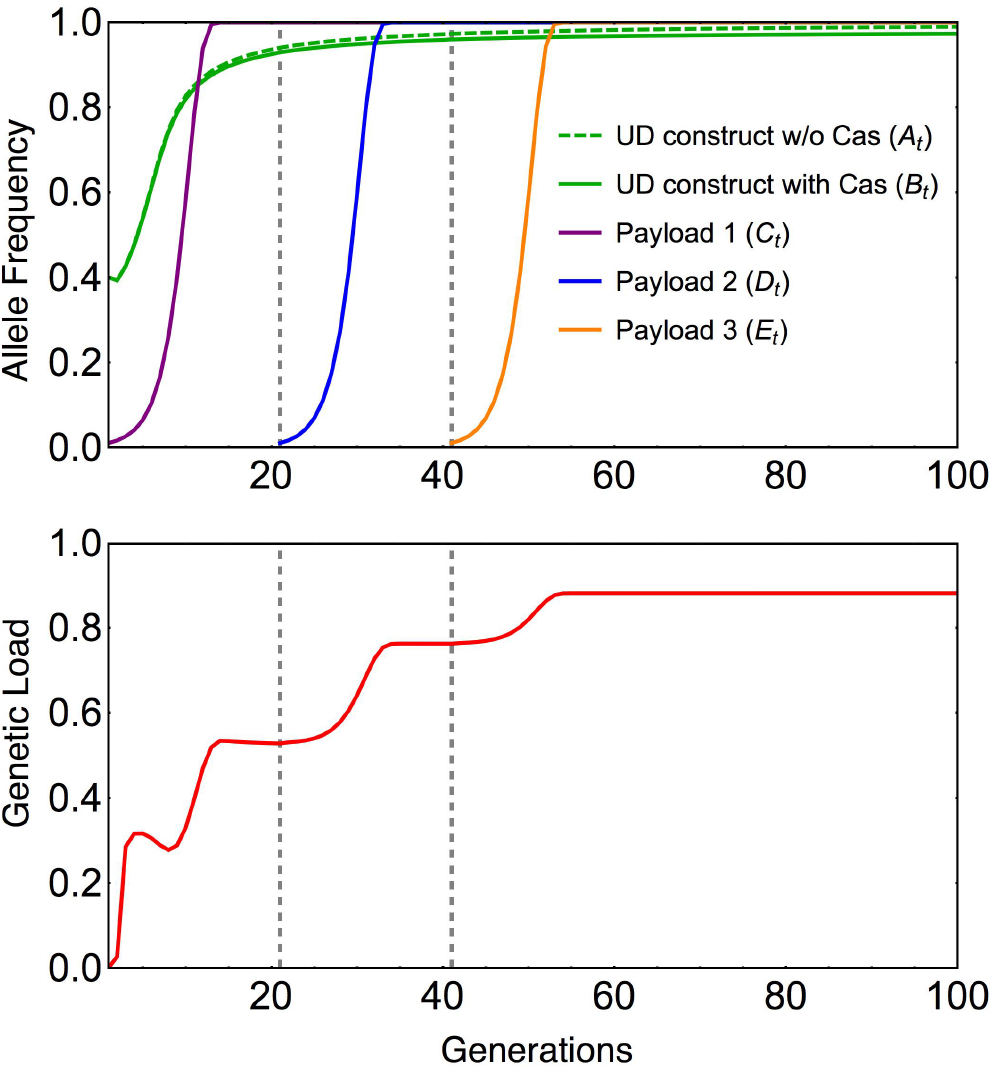
Time-series plots show frequencies of successively released homing constructs with new payloads (top panel) and the gradual buildup of genetic load in the population with the spread of each new payload gene. The first release has starting frequency of underdominance component at 40% and the first payload at 1%. Each successive payload is released to achieve 1% starting frequency. Each payload gene has a 50% female-limited homozygous fitness cost. Dashed grey lines show the time of release of successive payloads. Other parameters are *s_c_*=0.05, *H* = 0.95.

**Figure 6:**
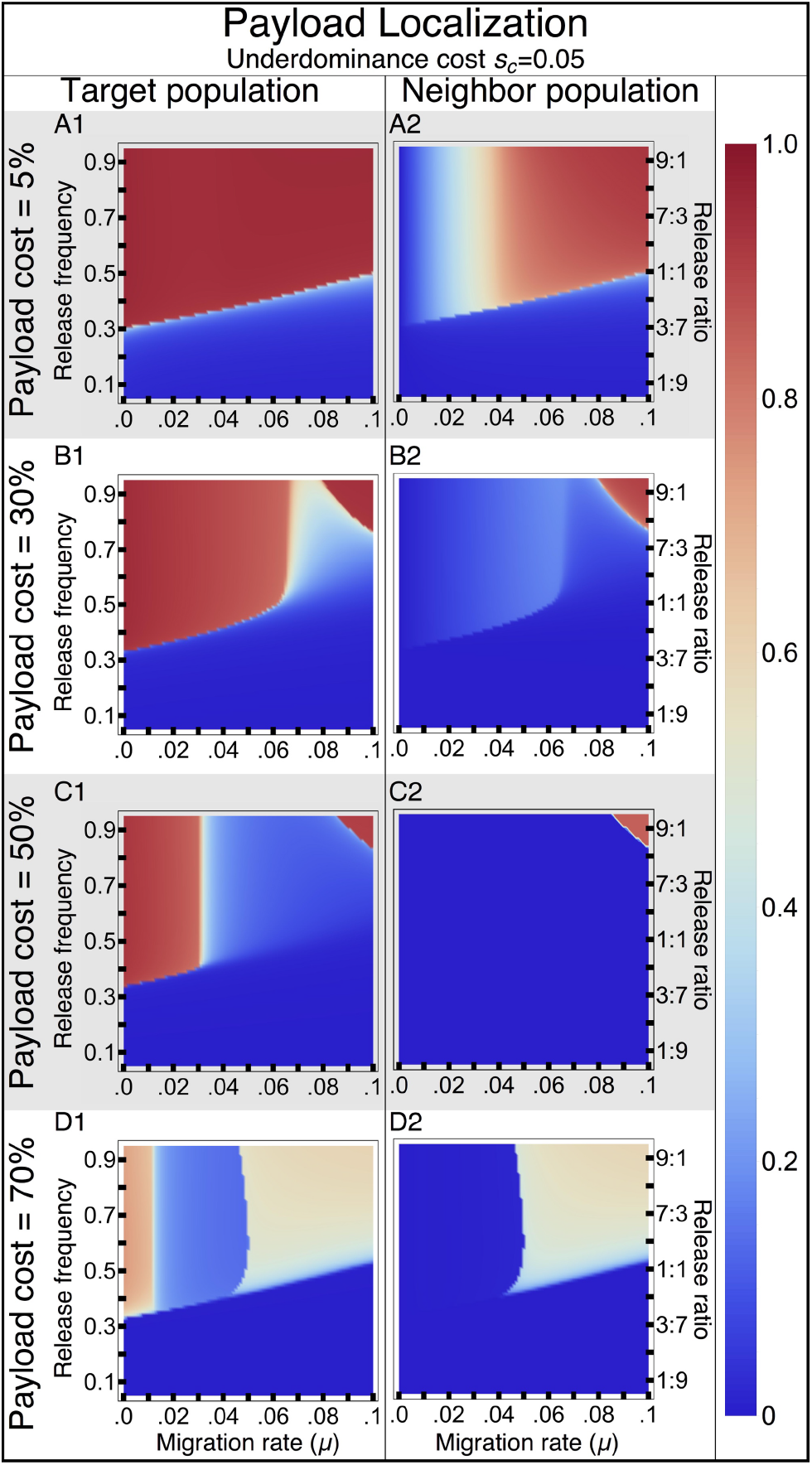
Localization of the UTH drive: Colors show mean frequency of the payload allele over 100 generations in the Target and a Neighbor population. Cost of the underdominance component, *s_c_* = 0.05. Different rows show UTH drives with different homozygous payload costs. The two-genotype drive is released to attain a starting homing construct frequency of 0.05. The starting frequency of all released individuals (including those with the full drive complement and those with only the underdominance component) is given on the vertical axis in each panel. Payload costs described are for homozygotes of both sexes.

When payload genes affect fitness of both sexes, sequentially driving multiple payload genes that each have a small effect on fitness can achieve a much higher combined genetic load compared what can be achieved by driving a single high-cost payload that affects both sexes (supplementary figure S4).

### Localization of the gene drive

Similar to other gene drives with introduction thresholds, the influx of wild-type individuals into the target population with increasing migration rate increases the threshold frequency for the UTH drive (Figure 6). The level of localization of the drive depends upon the fitness costs associated with its two components. A UTH drive with a low-cost underdominance component (*s_c_*=0.05) fails to remain localized when driving a payload gene designed for population replacement (*s_p_*=0.05), unless migration rates are <1% (Figure 6A and Figure 7A,B). However, the UTH drive can achieve localized population alteration with high cost payload genes over a much broader range of migration rates compared to simple one-locus engineered underdominance drive or the daisy-chain drive (Figure 6B-D and Figure 7C,D,E).

**Figure 7:**
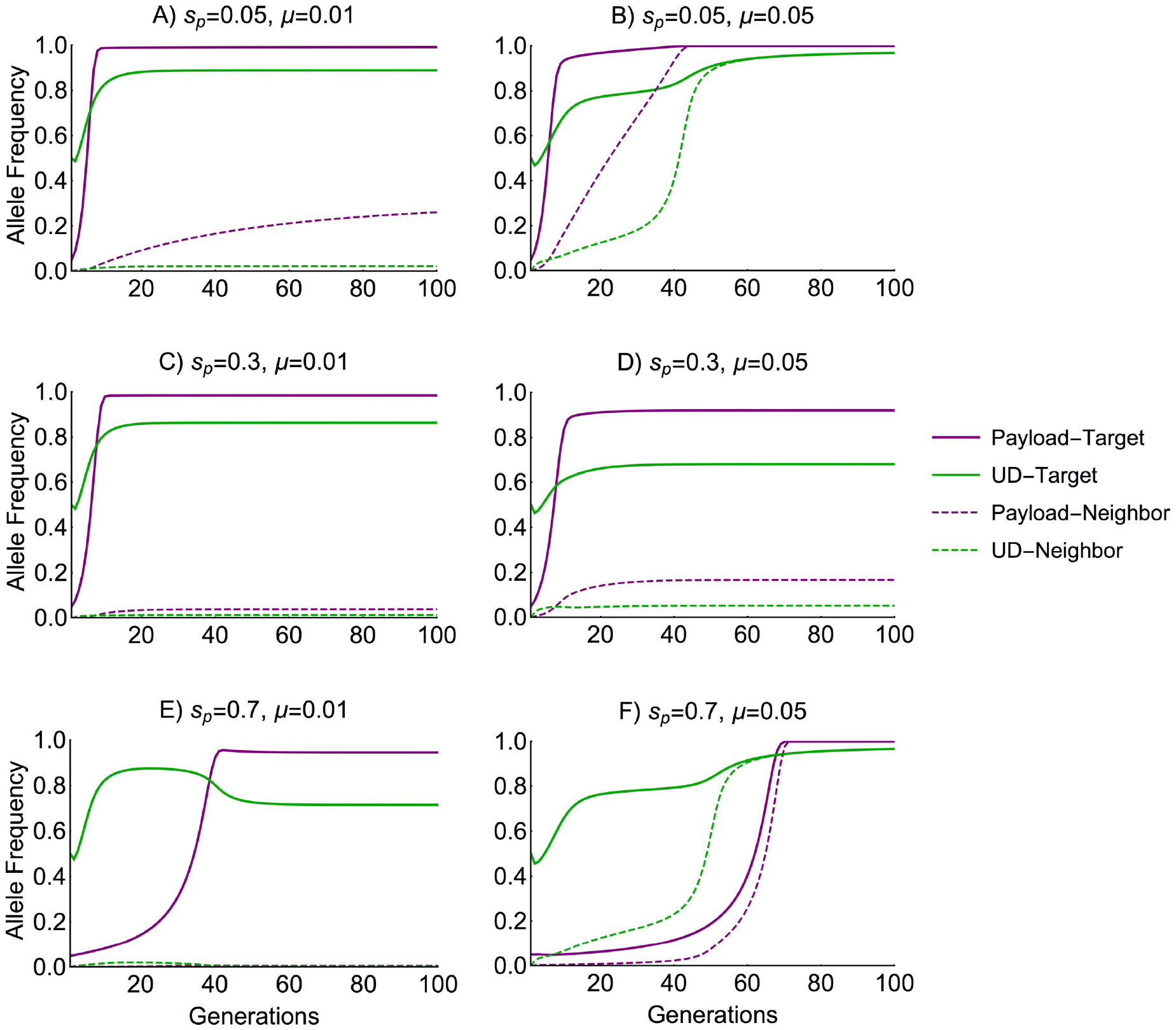
Time-series plots show the allelic frequencies of the payload (allele C_*t*_, purple lines) and the underdominance with Cas endonuclease (allele B_*t*_, green lines) in the target (solid lines) and the neighboring (dashed lines) populations. Individual plots show dynamics with low (*μ*=0.01, left column) or high (*μ*=0.05, right column) migration rates, and with different payload costs (across rows). Dynamics of the underdominance construct without Cas (allele At) are similar to that of allele B_*t*_, and are not shown for visual clarity.

The level of localization of a UTH drive can be greatly improved with certain modifications. For example, a UTH drive with underdominance cost (*s_c_*) of 0.2 has localization level much more similar to that of a simple two-locus engineered underdominance drive when driving payload genes of higher cost (supplementary figure S6; see Dhole et al. 2018). A modified, though perhaps more complex, design for a UTH drive could include a third toxin gene on the homing construct, that is suppressed by one of the suppressors on the underdominance component (Figure S7). The toxin gene on the homing construct from this modified UTH drive prevents it from diffusing into a neighboring population without the underdominance constructs, and therefore this modified UTH drive displays much higher level of localization (Figure S8).

For a small range of extremely high payload costs (Figure 6D and Figure 7E,F), the UTH drive with low-cost underdominance component can actually become less localized than a drive with lower cost payloads. This is because the extremely high payload costs keep the frequency of the homing construct (which carries the payload) at low levels until underdominance (which here confers only 5% fitness cost) becomes established in both populations, subsequently allowing the payload to spread in both populations (Figure 7F). A UTH drive with higher cost of underdominance is less likely to exhibit this behavior (Figures S7 and S8).

## DISCUSSION

Recently, spatially and temporally restricted gene drives have gained considerable attention as having less risk compared to unrestricted drives (e.g. NASEM 2016). The focus of most of the concern has been related to risk of unrestricted gene drives aimed at population suppression or eradication. A number of strategies for spatially restricted gene drives have been outlined (Davis et al. 2001, Marshall and Hay 2012a, Akbari et al. 2014, Buchman et al. 2018, Rasgon 2009, Noble et al. 2016, Burt and Deredec 2018) and a few have been tested on *Drosophila* in the lab (Akbari et al. 2013, Akbari et al. 2014, Reeves et al. 2014, Buchman et al. 2018), but mathematical models indicate that while these approaches are reasonable for changing characteristics of local populations (i.e. population replacement), they are less likely to be effective in suppressing local pest populations that have realistic density-dependent dynamics (but see Marshall and Hay 2014, Khamis et al 2018).

In general, it is difficult to effectively add high cost payloads to gene drives that intrinsically have high release thresholds (Marshal and Hay 2012a, Dhole et al. 2018). Unfortunately, gene drives with low intrinsic thresholds are less likely to remain localized. The work outlined here was aimed at developing an approach that would enable addition of substantial fitness cost to one spatially restricted drive strategy, two-locus underdominance, while maintaining an attainable release threshold.

The localization level of a TH drive is highly dependent on the drive used as anchor. The dynamics shown here are specific to a TH system based on two-locus engineered underdominance. Moreover, our basic proof-of-principle model ignores many biological factors that can influence the spread and localization of gene drives in real populations (Altrock et al. 2010, Huang et al. 2011, Edgington and Alphey 2017, Champer et al. 2018a). But the outcomes from this model are encouraging, especially in cases of invasive populations on oceanic islands where migration rates from the target population to a neighboring population are low. Even if the payload-associated fitness cost is only 30% and the neighboring population is initially the same size as the target population, spread through the neighboring population is not expected under these conditions (figure 6). One example that fits this category is rodents (i.e. mice and rats) that have invaded oceanic islands and are contributing to extinction of native flora and fauna (Bellard et al. 2016). Another example is the mosquito, *Culex quinquefasciatus*, that transmits bird malaria to the endangered Hawaiian honeycreepers (van Riper et al. 1986, Atkinson et al. 1995, Woodworth et al. 2005). For these cases it is useful to compare results for the UTH drive in figure 6 to results in Dhole et al. (2018) for two-locus underdominance. For payloads with equal impacts on male and female fitness, it is feasible to establish payloads with higher fitness cost when using UTH instead of the two-locus underdominance alone, and the release thresholds are substantially lower especially at higher payload fitness costs. For example, a UTH drive can establish a payload with 50% fitness cost with a much smaller release and under much higher migration rates compared to a two-locus underdominance drive (compare figure 6C here with figure 5h in Dhole et al. 2018). If the payload only has an impact on female fitness (figure 4), then the genetic load from the UTH on a local population can reach above 0.9 within 20 generations (i.e. only a few years for mice and mosquitoes on tropical islands).

Density dependence, as well as other factors such as mating behavior, stochasticity, spatial dynamics can give rise to complex population dynamics in response to perturbations. Therefore, the relationship between a specific amount of genetic load and the impact it has on population density or persistence is not straight forward. Recent analyses (Edgington and Alphey 2018, Khamis et al. 2018) also highlight that the specific values of several ecological parameters can have a large influence on the success of a gene drive. One advantage of the UTH system is that once one payload is established in a population, it is possible to add successive payloads (figure 5) by releasing a small number of additional individuals. One could start with a payload that results in a relatively low genetic load, and that load could be increased as needed. In cases where payload transgenes with female-limited fitness effects are not easily available, the ability of UTH drive to successively drive multiple payloads can be used to achieve higher genetic loads than are possible with single payload genes that affect both sexes (Figure S4).

It is important to recognize that a genetic load that decreases density of the invasive pest but does not cause eradication could be more sustainable than a high load that caused local extinction because the UTH with moderate load would remain active in the target population and would therefore impact any rare immigrants arriving from other populations. The ability to adjust the genetic load would be helpful in these cases where the goal was just to lower the population density below a harmful level.

One goal for localized gene drives is to prevent the spread of engineered genes across international boundaries. However, such demarcations often do not fall along ecological barriers to gene flow. In such scenarios, migration rates between populations may be too high (greater than 3% per generation) to ensure spatial restriction with a UTH drive. Curiously, when the payload cost is over 70% there is also some risk of impact on the neighboring population (figure 6). The reason for this is that due to the high fitness costs and high gene flow, the homing construct that contains the payload remains at low frequency for many generations. This allows the underdominance component to become established in both populations, then driving the payload to high frequency in both populations (Figure 7F).

The level of localization of a UTH drive could be improved by designing the underdominance component to have a higher fitness cost, or by including a toxin gene on the homing construct that is suppressed in the presence of the underdominance constructs (Figure S7). The third toxin would impose a genetic background-dependent cost on the homing construct, so that it can spread in the target population largely unimpeded by the toxin, but be eliminated quickly from the neighboring populations that have very low levels of the underdominance constructs.

The feasibility of these systems as useable tools would depend on the practical difficulties involved in engineering these components as well as their effects on fitness in natural conditions. Localized alteration of natural populations may require combining multiple approaches. For example, a TH drive with a homing construct that targets an endemic/private or locally fixed wild-type allele (Sudweeks et al. in review) is likely to be much more localized than a TH drive that targets alleles present at high levels in multiple populations. The localization of a gene drive designed for population suppression could also be improved if the payload gene that reduces fitness also reduces the likelihood of migration of engineered individuals, for example, by reducing mobility or tolerance to the elements during migration.

All of the results presented here come from a general model designed to introduce the concept of tethered homing drives, and should not be overinterpreted as predicting an outcome in a specific case or immediate suitability for use in real populations. The analyses shown are intended to facilitate a comparison between the general behaviors of the UTH drive with previously proposed gene drives. More detailed models that reflect the biology of a targeted population of a species will be needed for assessing whether the UTH drive or related TH drives are appropriate for a given problem.

## Acknowledgements

We are grateful to Jennifer Baltzegar, Brandon Hollingsworth, Alex DeYonke, Jaye Sudweeks and Michael Vella for helpful discussion and comments on previous versions of the manuscript. SD was supported by a grant from the W. M. Keck Foundation to FG. ALL were funded by National Institute of Health grant R01-AI091980 and a National Science foundation grant RTG/DMS—1246991 to FG.

